# Stabilizing proteins through saturation suppressor mutagenesis

**DOI:** 10.1101/2021.08.07.455542

**Authors:** Shahbaz Ahmed, Kavyashree Manjunath, Gopinath Chattopadhyay, Raghavan Varadarajan

## Abstract

While there have been recent, transformative advances in the area of protein structure prediction, prediction of point mutations that improve protein stability remains challenging. It is possible to construct and screen large mutant libraries for improved activity or ligand binding, however reliable screens for mutants that improve protein stability do not exist, especially for proteins that are well folded and relatively stable. We demonstrate that incorporation of a single, specific destabilizing, (parent inactive) mutation into each member of a deep mutational scanning library followed by screening for suppressors, allows for robust and accurate identification of stabilizing mutations. When coupled to FACS sorting of a yeast surface display library of the bacterial toxin CcdB, followed by deep sequencing of sorted populations, multiple stabilizing mutations could be identified after a single round of sorting. Multiple libraries with different parent inactive mutations could be pooled and simultaneously screened to further enhance the accuracy of identification of stabilizing mutations. Individual stabilizing mutations could be combined to result in a multimutant with increase in thermal melting temperature of about 20 °C and enhanced tolerance to high temperature exposure. The method employs small library sizes and can be readily extended to other display and screening formats to rapidly isolate stabilized protein mutants.

## Introduction

Directed evolution has drastically reduced the time required to engineer desired function into proteins (1–4). Enzymes and other proteins with altered function or binding specificity have been evolved using yeast surface display (YSD), phage display (5, 6) or other *in vivo* functional screens (7–9). Phage display utilizes its surface proteins pIII and pVIII, which are fused to the protein of interest (10). Phage display can be used to generate libraries of very high diversity (11) which can be screened for binding to a target ligand. Agglutinin based Aga2p is a widely used system to display proteins on the yeast cell surface (5). Aga2p is a small protein, covalently linked via disulphide linkages to the yeast cell surface protein Aga1(12). Different populations in a yeast library can be enriched using FACS. Relative to the phage display library, sizes are lower (∼10^11^ vs 10^7^). However, with YSD, eukaryotic post translation modifications are possible. While screening for mutants with improved binding or enzymatic activity is straight forward, it is non-trivial to screen for mutants with improved stability. Deep mutational scanning is an approach which combines screening/selection of a mutant library with next generation sequencing to identify the degree of enrichment of mutants following a selection or screen, relative to the population present in the original library (13–16). Some prior studies have suggested that stabilized mutants are expressed at higher levels than wild type (WT), however, several other studies have not observed this (17–19). In several cases it was observed that for stable proteins, mutants with improved stability are expressed at a similar level to WT (18–21) and hence expression alone cannot be used to discriminate mutants with higher stability from mutants with a slightly destabilized phenotype. We recently showed that the amount of active protein on the yeast cell surface (detected by the amount of bound ligand) correlates better with *in vitro* thermal stability or *in vivo* solubility than the amount of total protein on the yeast cell surface, for destabilized mutants (22). However, mutants above a certain stability threshold show similar expression and ligand binding to wild type irrespective of their stability. Previously, to find stabilized variants of proteins with high intrinsic stability, YSD libraries were subjected to thermal stress, to enrich for more stable variants followed by sorting to identify variants which retained binding to a conformation specific ligand (23, 24). While this is potentially useful, yeast cells cannot replicate after high temperature exposure, and hence the method requires repeated rounds of plasmid isolation, PCR amplification and retransformation in yeast cells after each round of enrichment. Also, if a protein exhibits reversible thermal unfolding, enrichment of stabilized mutants in such cases will be difficult. An alternative approach to isolate stabilizing mutations is to introduce a destabilizing mutation, hereafter referred to as parent inactivation mutation (PIM), and then create mutant libraries in this background to screen for suppressors (25–31). Often, this methodology requires multiple rounds of enrichment to isolate stable mutants. The reversion of PIM to wild type or non-destabilizing mutants during library generation or enrichment enriches for mutants lacking the PIM instead of desired PIM-suppressor pairs. Additionally, in multi-round format, this methodology potentially allows isolation of only a few stabilizing mutations, and does not distinguish between allele specific and global suppressors. It is also unclear whether it will always be possible to isolate suppressors for every PIM. In the present study, we have modified this approach by introducing PIMs in the background of a deep mutational scanning library of bacterial toxin CcdB, sorted different populations, subjected each population to deep sequencing of the ccdb gene and reconstructed the mean fluorescent intensity (MFI) of each mutant as described (22) (see also Experimental Procedures section). We use both the reconstructed binding MFI (MFI_seq_ (bind)) and expression MFI (MFI_seq_ (expr)) as criteria to differentiate between stabilized mutants, WT-like and destabilized mutants. This single-site, saturation suppressor mutagenesis (SSSM) methodology (Figure 1), was described previously (29). Using this methodology, two different types of suppressors can be identified. Proximal suppressors reverse the destabilizing effect of the PIM by locally compensating packing defects caused by the PIM. Proximal suppressors are allele specific and do not show stabilizing effects as a single mutant in the absence of the PIM (29, 32). Distal suppressors are located far from the PIM and often act as global suppressors, reversing the effects of multiple individual PIMs. A distal suppressor also typically stabilizes the protein relative to WT, in the absence of the PIM. Using this methodology, we could readily identify putative stabilized mutants with a minimal number of false positives.

**Figure 1:**
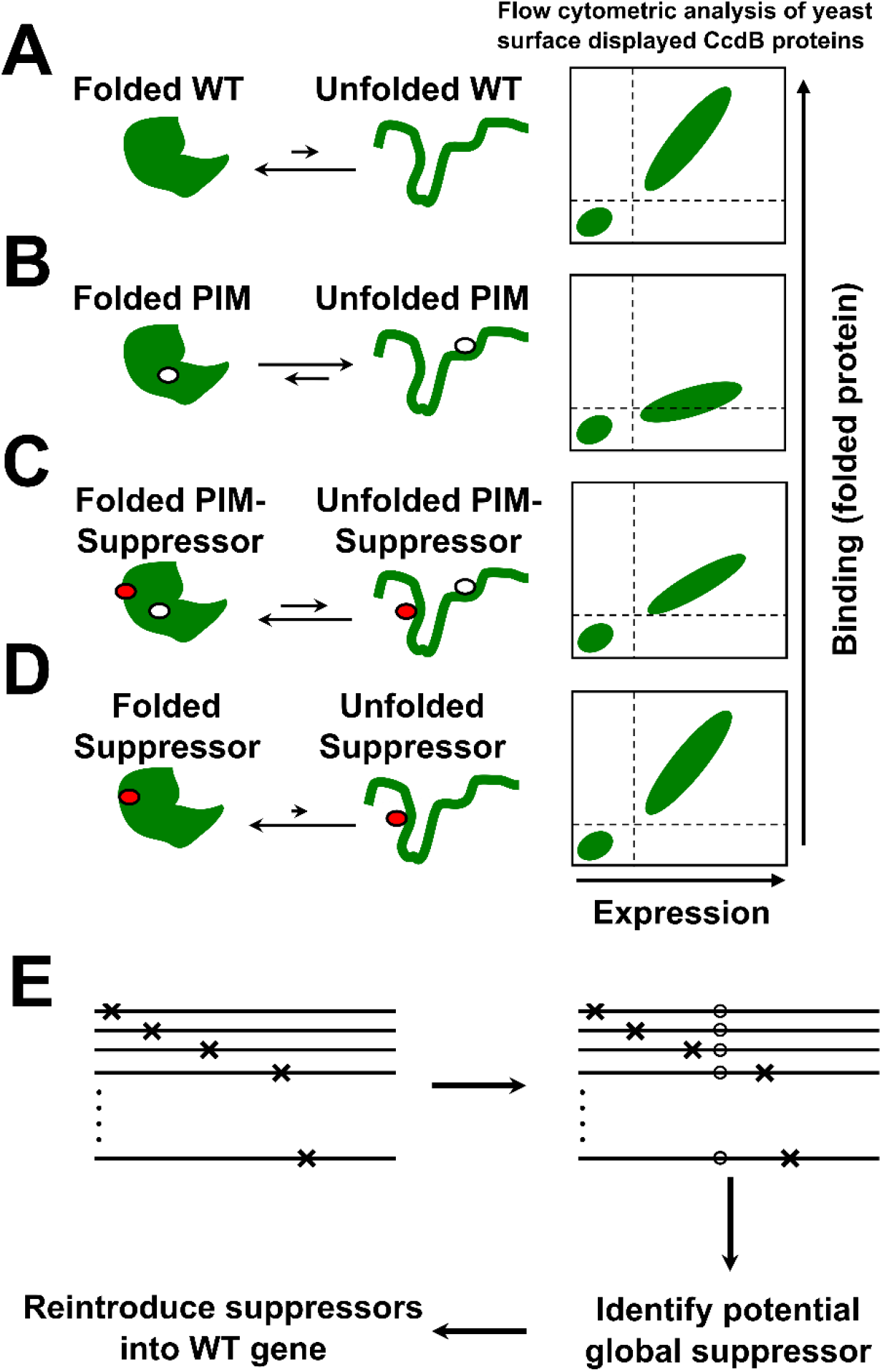
Schematic representation of Second Site Suppressor Mutagenesis (SSSM) methodology. Proteins exist in an equilibrium between folded and unfolded states. (A) WT proteins generally have the equilibrium shifted towards the folded state. Such proteins when expressed on the yeast cell surface show good expression and binding to their cognate ligand (B) Introduction of a <u>P</u>arent Inactive <u>M</u>utation (PIM) shift the equilibrium towards the unfolded state, and decrease the amount of properly folded protein on the yeast cell surface, leading to decreased ligand binding. (C) Second-site suppressor mutation, distal from and present in the background of the PIM will reduce the amount of unfolded protein present at equilibrium. Such double mutants have higher expression and binding on the yeast cell surface compared to the PIM alone. (D) Such global/distal suppressor mutations can stabilize multiple PIMs and also stabilize the WT protein, although the expression and binding of suppressor alone on the yeast cell surface is similar to the WT protein. (E) Saturation suppressor libraries are created by introducing a PIM (o) into the background of a deep mutational scanning library, selecting for suppressors, and reintroducing identified suppressors in the background of the WT gene.

## Results

### Selection of PIM and sorting of Second Site Saturation Mutagenesis (SSSM) library of CcdB

CcdB is a bacterial toxin that causes bacterial cell death by binding and poisoning DNA gyrase (33). When expressed on the surface of yeast, properly folded CcdB can be detected by binding to FLAG tagged GyrA14 followed by incubations of cells with anti FLAG primary and fluorescently labeled secondary antibody (29). We selected four PIMs based on their *in vitro* thermal stability and *in vivo* activity in *E*.*coli* (14, 29). PIM V18D is completely inactive and highly aggregation prone due to a charged mutation in the core of the protein. V18G and V20G are partially inactive *in vitro* due to the formation of a cavity in the core of the protein, with V20G showing higher activity *in vivo* compared to V18G (14). L36A is the most active amongst the four PIMs, the mutant protein is partially aggregated (29). SSSM libraries were constructed by introducing each PIM individually in the background of the deep mutational scanning library (29). These PIMs and their corresponding SSSM libraries showed variable expression and binding depending on the PIM present in the single mutant library (Supporting Figure S1). In each library, binding and expression experiments had slightly different numbers of mutants for which MFI_seq_ was calculated (Supporting Table S1). Different populations of SSSM libraries were sorted based on the expression and binding histograms of the libraries (Supporting Figure S2). Populations were subjected to deep sequencing and the mean fluorescence intensities of binding (MFI_seq_ (bind)) and expression (MFI_seq_ (expr)) were estimated as described (22). Values of normalized MFI_seq_ (bind) and normalized MFI_seq_ (expr) were used to predict stabilized mutants. We only considered mutants which were present in more than one library. Putative stabilized mutants were identified as those which have a normalized MFI_seq_ value for either binding or expression >1.25. WT like mutants were those which had normalized MFI_seq_ values of 0.9-1.25 and destabilized mutants were those which had normalized MFI_seq_ values of <0.9. Within each predicted class, mutants were randomly selected for experimental validation. 112 individual point mutants in the WT background were expressed and purified and the T_m_ was measured using a thermal shift assay (TSA) (Supporting Table S2). We found a better correlation between the MFI_seq_ (bind) and the stability of stabilized and marginally destabilized (0>ΔT_m_>-5) mutants compared to MFI_seq_ (expr) for all the libraries (Figure 2A-H). In the case of V18D library, the overall correlation between MFI_seq_ values of expression or binding with stability of the mutants was poor (Figure 2A, 2E), nevertheless the best binders showed significant stabilization. The remaining libraries showed a good correlation of stability with MFI_seq_ (bind) or MFI_seq_ (expr), MFI_seq_ (bind) showed a better correlation than MFI_seq_ (expr) in all the cases.

**Figure 2:**
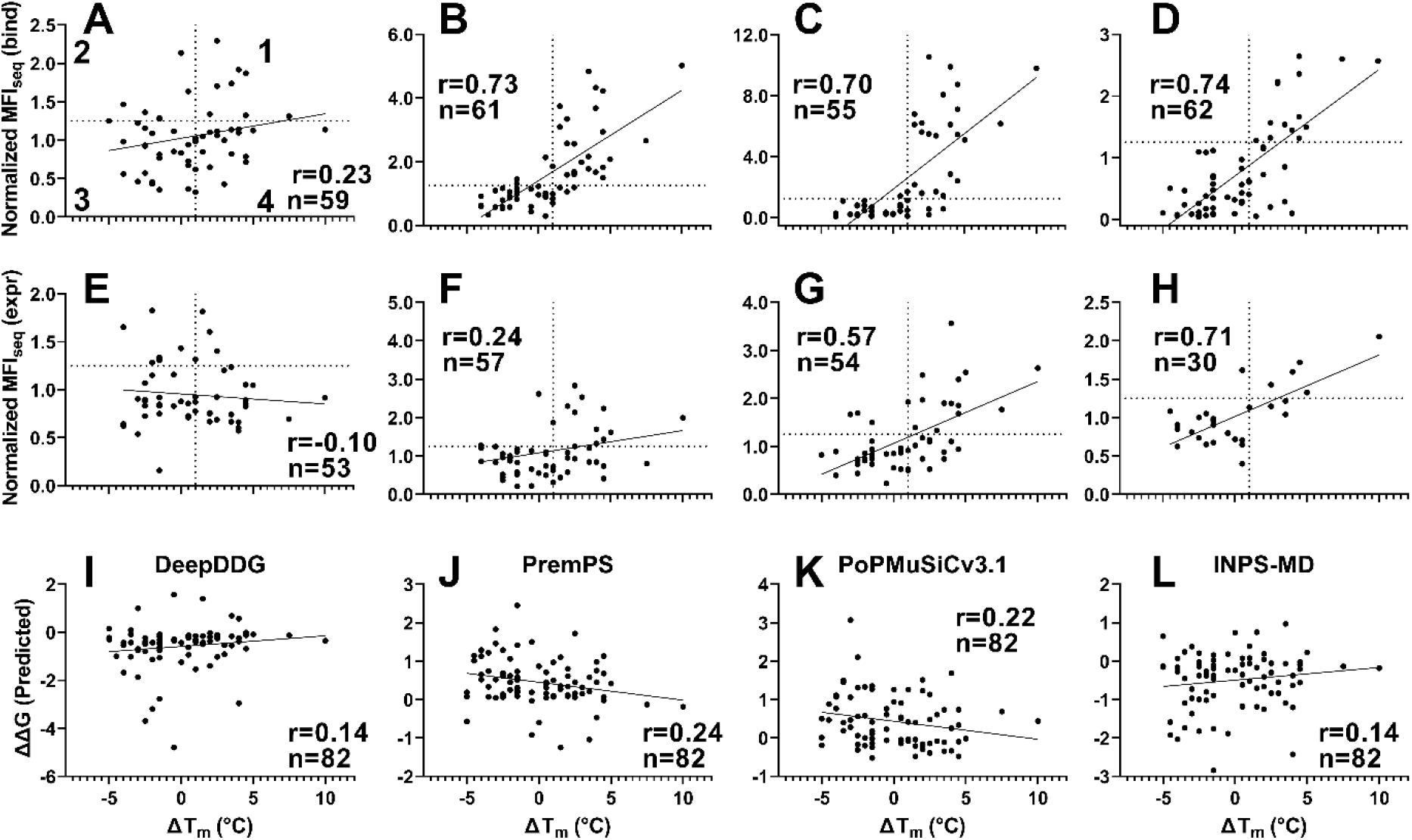
Correlation of CcdB mutant stability (ΔT_m_ of single mutant) with normalized MFI_seq_ (bind) or MFI_seq_ (expr) of (PIM, mutant) pair in PIM libraries. MFI_seq_ values of double mutants were normalized with the MFI_seq_ values of their respective PIM. First, second, third and fourth quadrants numbered in (A) represent true positive, false positive, true negative and false negative points respectively. Normalized MFI_seq_ (bind) correlation with thermal stability for V18D (A), V18G (B), V20G (C) and L36A (D) libraries. Normalized MFI_seq_ (expr) correlation with thermal stability for V18D (E), V18G (F), V20G (G) and L36A (H) libraries. Normalized MFI_seq_ (bind) correlates better than Normalized MFI_seq_ (expr) with thermal stability. For V18D while overall correlation is poor, those mutants with highest MFI_seq_ (bind) are stabilized. Thermal stability predictions by *in silico* methods (I) DeepDDG, (J) PremPS, (K) PoPMuSiCv3.1 and (L) INPS-MD. For *in silico* stability measurents, mutants which were present in any libaray were used. Predicted ΔΔG >0 is stabilizing in the case of DeepDDG and INPS-MD, Predicted ΔΔG <0 is stabilizing in the case of PremPS and PoPMuSiC.

In a previous report, we compared the accuracy of predictions from several *in silico* tools with our experimental method to estimate the relative stability for several destabilized mutants of CcdB (22). Four *in silico* tools, DeepDDG (34), PremPS (35), PopMuSiC (36) and INPS-MD (37) which calculated the ΔΔG of mutant (ΔG of unfolding of mutant-ΔG of unfolding of WT) of CcdB mutants showed a correlation of greater than 0.5 with the corresponding *in vitro* measured thermal stability. However, in the case of marginally destabilized and stabilized mutants examined in the present work, the predictions from these programs showed very poor correlation with experimentally thermal stability (Figure 2I-L).

In related work, we have stabilized the receptor binding domain of the Spike S protein of SARS-COV-2 using saturation suppressor mutagenesis (38). We identified several stabilizing mutations and could stabilize the protein by 8 °C. We used PROSS to predict stabilizing mutations in RBD. PROSS provided only two designs. Design 1 contained the S375D mutation and Design 2 contained S375D and V362I, from analysis of corresponding YSD data, both of these variants are predicted to have stabilities similar to or lower than WT (39). The poor performance of PROSS in this case is probably because of the lack of sufficiently diverse sequence data to identify stabilizing mutations. In the case of CcdB, PROSS predicted several mutations to be stabilizing, however some were likely to be destabilizing according to our YSD data. Several predictions could not be validated as they wereeither part of the active-site or we did not have corresponding YSD binding data for the mutant. The PROSS results for CcdB are now summarized in Supporting Table S3.

In case of the V18D library, we observed that most mutations have low values of normalized MFI_seq_ (bind) as well as MFI_seq_ (expr). This might be because highly destabilizing PIM cannot be rescued by a single suppressor mutation. We observed a low sensitivity and specificity but reasonable accuracy of prediction for stabilized mutants when predictions were made based on either MFI_seq_ (bind) or MFI_seq_ (expr) (Figure 3, Table 1) for this library. Since the results were not as promising as the other libraries, the V18D library was not included in subsequent analyses. A similar analysis was performed for the other three libraries and we found a high sensitivity of prediction for stabilized mutants for these libraries when MFI_seq_ (bind) was used as the criterion (Table 1).

**Figure 3:**
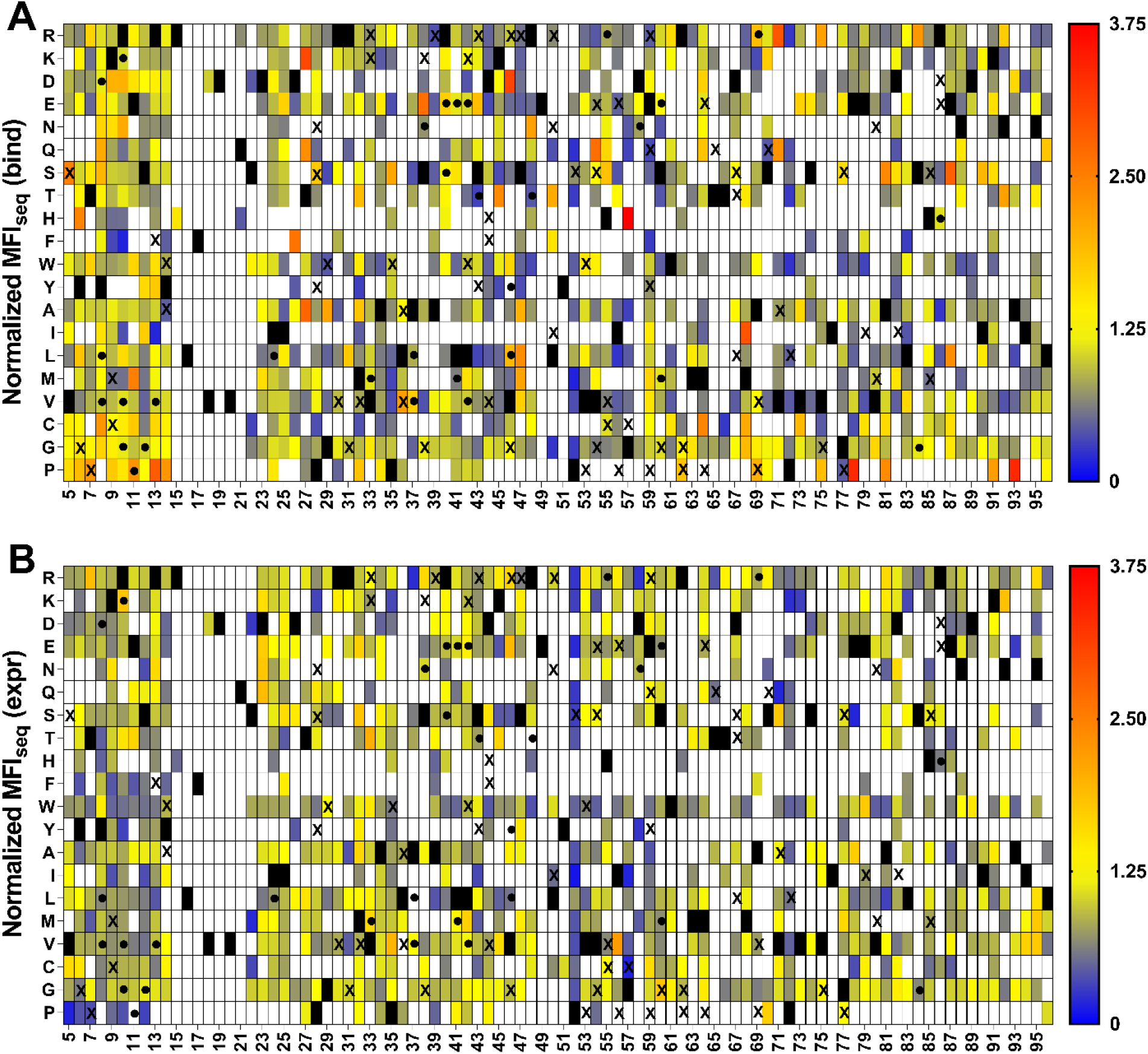
Heat maps of normalized MFI_seq_ (bind) and MFI_seq_ (expr) for V18D library. Normalized (with respect to V18D) values of MFI_seq_ (bind) (A) and MFI_seq_ (expr) (B) were categorized in different ranges. Mutants with normalized MFI_seq_ (bind) or MFI_seq_ (expr) >1.25 were categorized as putative stabilized mutants. Black rectangle represents WT residue, mutants with no data available are indicated with a white rectangle. Several single mutants were characterized *in vitro* to estimate their stability, stabilized mutants having ΔT_m_>1 are indicated with a “•” and destabilized mutants ΔT_m_<0 are indicated with an “X”. Mutants with normalized MFI_seq_ value ≥ 3.75 are coloured in red.

**Table 1:** Identification of stabilized mutant solely from normalized MFI_seq_ (bind) or MFI_seq_ (expr) data. Mutants were predicted to be stabilized if the corresponding MFI_seq_ value was greater than 25 percent. CcdB mutant thermal stability data was taken from previous studies (22, 54) as well as additional mutant stability data measured in this study (Supplementary Table S2).

| Parameter | V18D library |  | V18G library |  | V20G library |  | L36A library |  |
| --- | --- | --- | --- | --- | --- | --- | --- | --- |
|  | Bind | Expr | Bind | Expr | Bind | Expr | Bind | Expr |
| **Sensitivity<sup>a</sup>** | 0.27 | 0.15 | 0.80 | 0.44 | 0.79 | 0.54 | 0.63 | 0.56 |
| **Specificity<sup>a</sup>** | 0.73 | 0.88 | 0.89 | 0.79 | 0.98 | 0.78 | 1.00 | 0.91 |
| **Accuracy<sup>a</sup>** | 0.57 | 0.63 | 0.86 | 0.67 | 0.90 | 0.70 | 0.89 | 0.84 |

<sup>a</sup>The sensitivity, specificity and accuracy were calculated using the following formulae, where TP, TN, FP and FN correspond to true positive ( $\Delta T_m$ (predicted) > 0, $\Delta T_m$ (observed) > 0), true negative ( $\Delta T_m$ (predicted) < 0, $\Delta T_m$ (observed) < 0), false positive ( $\Delta T_m$ (predicted) > 0, $\Delta T_m$ (observed) < 0) and false negative ( $\Delta T_m$ (predicted) < 0, $\Delta T_m$ (observed) > 0) respectively..

SSSM libraries with V18G or V20G as PIM displayed the highest sensitivity of prediction of stabilizing mutations (Figure 4, Figure 5). We found some false positives for the L36A SSSM library (Figure 6), which had similar expression and binding compared to the deep mutational scanning library of WT CcdB (Supporting Figure S1). When stabilized mutant predictions were made based on the MFI_seq_ (expr), we observed lower sensitivity compared to the predictions based on MFI_seq_ (bind) (Table 1) for all libraries.

**Figure 4:**
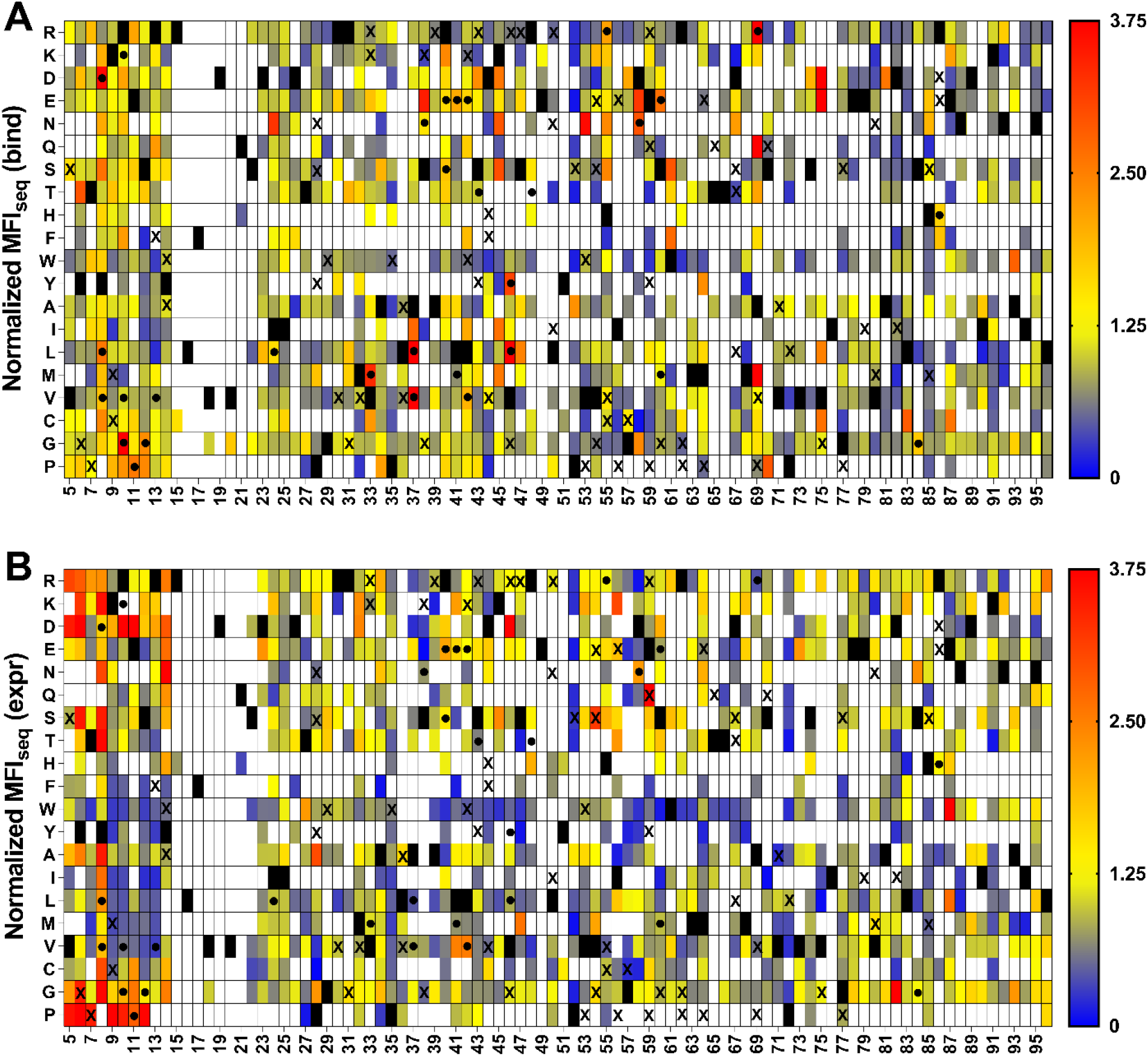
Heat maps of normalized MFI_seq_ (bind) and MFI_seq_ (expr) for V18G library. Normalized (with respect to V18G) values of MFI_seq_ (bind) (A) and MFI_seq_ (expr) (B) were categorized in different ranges. Mutants with normalized MFI_seq_ (bind) or MFI_seq_ (expr) >1.25 were categorized as putative stabilized mutants. Black rectangle represents WT residue, mutants with no data available are indicated with a white rectangle. Several single mutants were characterized *in vitro* to estimate their stability, stabilized mutants (ΔT_m_>1) are indicated with a “•” and destabilized mutants with (ΔT_m_<0) are indicated with an “X”. Mutants with normalized MFI_seq_ value ≥ 3.75 are coloured in red.

**Figure 5:**
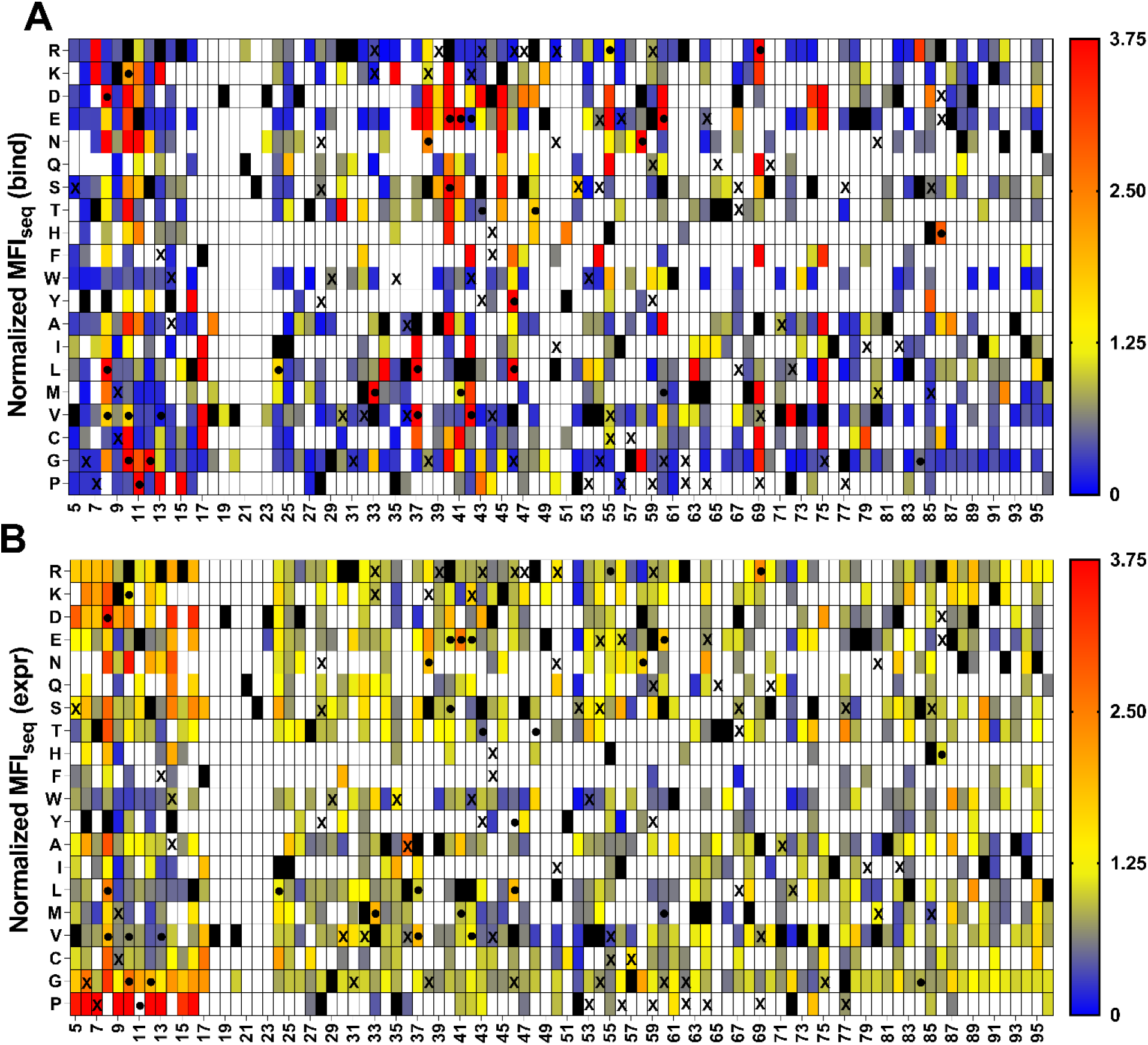
Heat map of normalized MFI_seq_ (bind) and normalized MFI_seq_ (expr) for V20G library. Normalized (with respect to V20G) values of MFI_seq_ (bind) (A) and MFI_seq_ (expr) (B) were coloured from blue to red as increasing MFI_seq_ values. Mutants with normalized MFI_seq_ (bind) or MFI_seq_ (expr) greater than 1.25 were categorized as putative stabilized mutants. Black rectangle represents WT residues, mutants where no data are available are indicated with a white rectangle. A subset of mutants were purified and their thermal stability (T_m_) *in vitro* was measured. Stable mutants (ΔT_m_>1) are indicated with a “•” and destabilized mutants (ΔT_m_<0) are indicated with an “X”. Mutants with normalized MFI_seq_ value ≥ 3.75 are coloured in red. It is clear that MFI_seq_ (bind) is superior to MFI_seq_ (expr) in identification of stabilized mutants.

**Figure 6:**
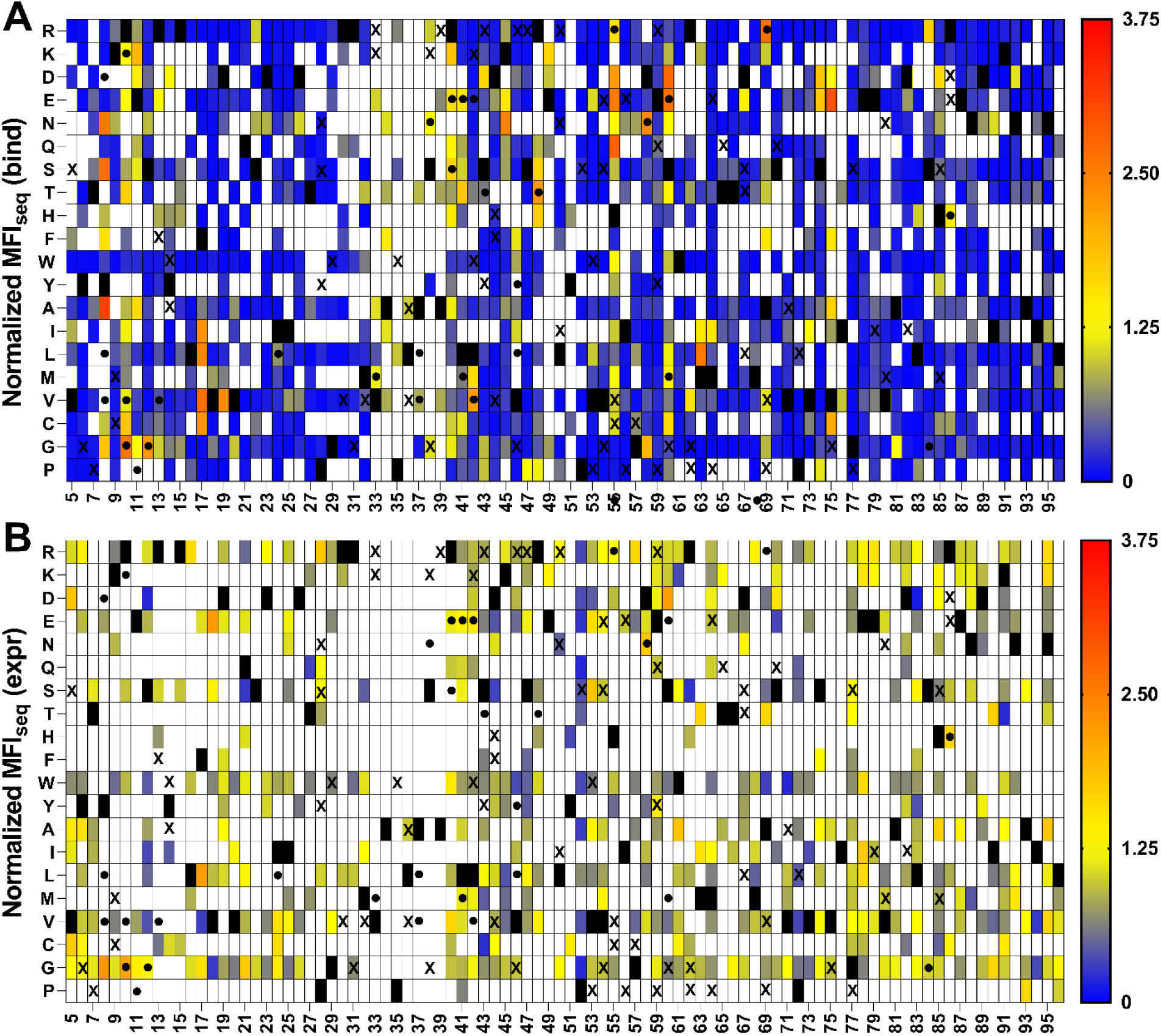
Heat maps of normalized MFI_seq_ (bind) and MFI_seq_ (expr) for L36A library. Normalized (with respect to L36A) values of MFI_seq_ (bind) (A) and MFI_seq_ (expr) (B) were categorized in different ranges. Mutants with normalized MFI_seq_ (bind) or MFI_seq_ (expr) >1.25 were categorized as putative stabilized mutants. Black rectangle represents WT residue, mutants with no data available are indicated with a white rectangle. Several single mutants were characterized *in vitro* to estimate their stability, stabilized mutants (ΔT_m_>1) are indicated with a “•” and destabilized mutants with (ΔT_m_<0) are indicated with an “X”. Mutants with normalized MFI_seq_ value ≥ 3.75 are coloured in red.

We hypothesized that the second mutation which alleviates the destabilizing effect of a PIM may act as a global suppressor and can therefore alleviate the destabilizing effects of other PIMs. To confirm this, common suppressors were shortlisted which were present in different PIM libraries (Figure 7A). True global suppressors were identified as those which alleviated the destabilizing effect of at least two PIMs. Using this criterion to identify stabilized mutants we found a sensitivity of 1 for the prediction. We also found that at several positions, there were multiple mutants with stabilizing phenotypes, which indicates that the WT is not the most preferred in terms of stability, this information can also be used as a criterion to find the stabilizing mutation. Interestingly none of the suppressors were found at the residues directly involved in GyrA binding. In any case, we have recently shown that such active-site residues can be identified based on the pattern of MFI_seq_ (bind) and MFI_seq_ (expr) in deep mutational scanning libraries and removed from the set of putative global suppressors (22).

**Figure 7:**
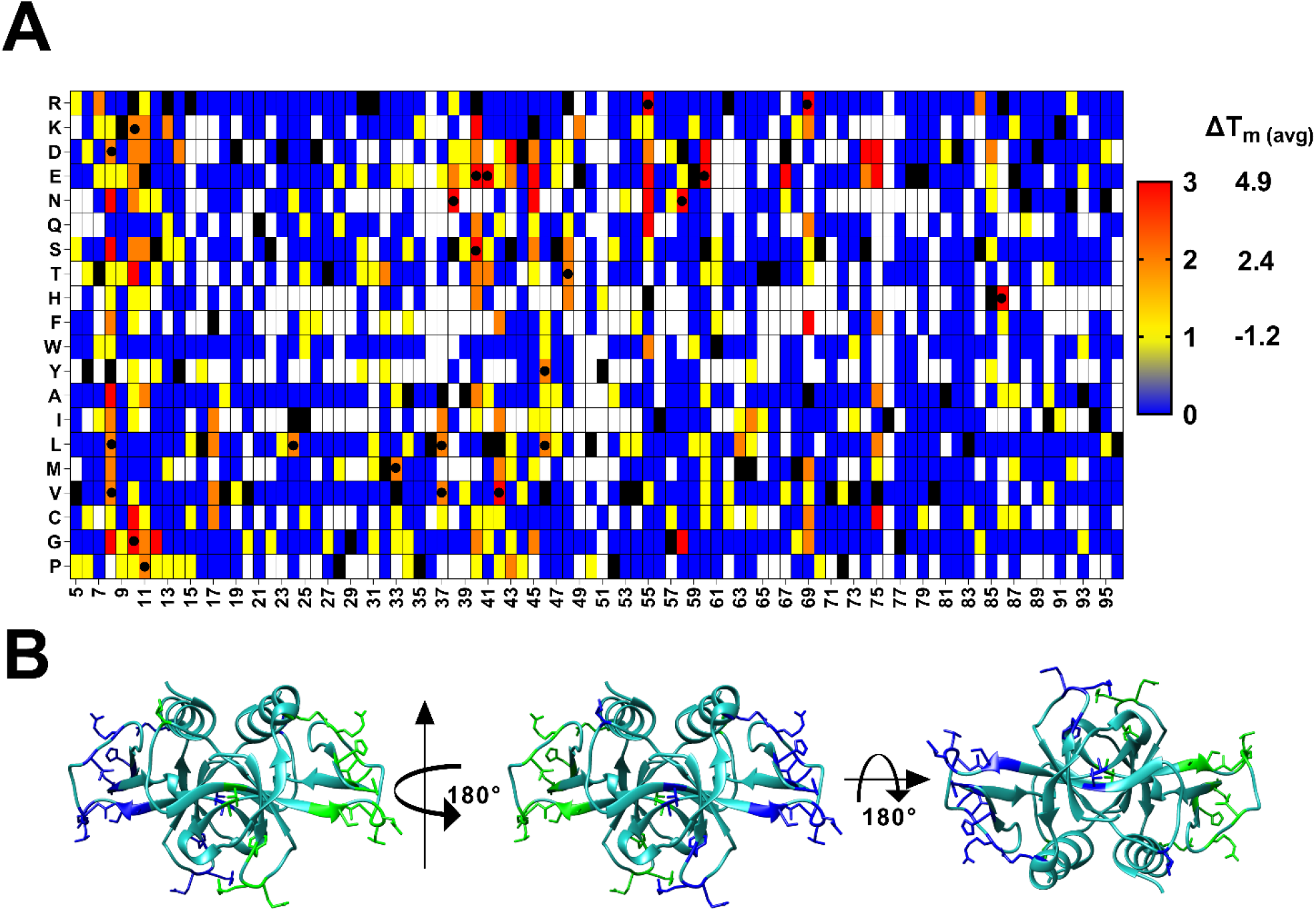
Putative stabilized mutants found in all the three V18G, V20G and L36A libraries based on MFI_seq_ (bind). (A) heat map of putative suppressor mutants in each library were given a score of one unit. Blue to red indicate that the mutant acted as a suppressor in zero, one, two and three different libraries. White indicate the mutants where no data are available. Experimentally confirmed stable mutants (ΔT_m_>1) are indicated with a “•” (B) The residue locations of experimentally characterized stabilized mutants are shown in blue and green colour for chain A and chain B of CcdB dimer respectively (PDB ID: 3VUB).

Most of the stabilizing mutations occur in surface exposed loop regions (Figure 7B) and a proper interpretation is only possible in the context of a high resolution structure. In another study, where we have carried out detailed mechanistic studies to understand the mechanistic basis for global suppressors in multiple protein systems, we have solved the structures of a few individual stabilizing mutations (S12G, V46L and S60E) identified in the present study (40). In the case of S12G, we found that two additional water molecules were present in the place of S12 and formed hydrogen bonds with the main chain of residue E11 and R13. In the case of V46L, a new hydrophobic interaction was introduced between side chains of M64 and V46L, as well as an additional hydrogen bond between the R62 side chain and main chain carbonyl group of residue 46. In the case of WT, a salt bridge is formed between R48 and E49, upon introduction of the S60E mutation, a loop flipping is observed and R48 now forms a new salt bridge with E60, which restricts the movement of the loop 41-50. It is not possible to anticipate these structural changes upon mutation in these mutants through modelling. We estimated the affinity of some of the stabilized and destabilized and multi-mutants of CcdB towards GyrA14 using SPR, and found that they retain affinities similar to WT CcdB, indicating that mutations at the non active-site positions do not alter the affinity of CcdB mutants for GyrA14 (Supporting Figure S3).

### Additive effect of stabilizing mutations

While single mutations usually do not enhance the stability of proteins to a great extent, individual stabilizing mutations can be combined resulting in multi-mutants with enhanced stability. We therefore constructed multiple double and triple mutants and measured their thermal stabilities using a thermal shift assay. For the generation of double and triple mutants, we used only a single criterion, namely that the centroid-centroid distance between any two of the mutants should be greater than 7 Å. All the double, triple and multi-mutants showed a higher T_m_ than the WT (Figure 8A). We also observed a good correlation (r =0.98) between ΔT_m_ of double/triple mutants with the sum of ΔT_m_ of individual mutants (Figure 8B). When more than 7 mutants were combined such additive effects were not observed. While combining 7 or more stabilizing mutations, we did not consider any centroid-centroid distance cut-off and combined mutants with highest stability, the lack of additivity here indicates possible epistatic interactions between residues in close proximity.

**Figure 8:**
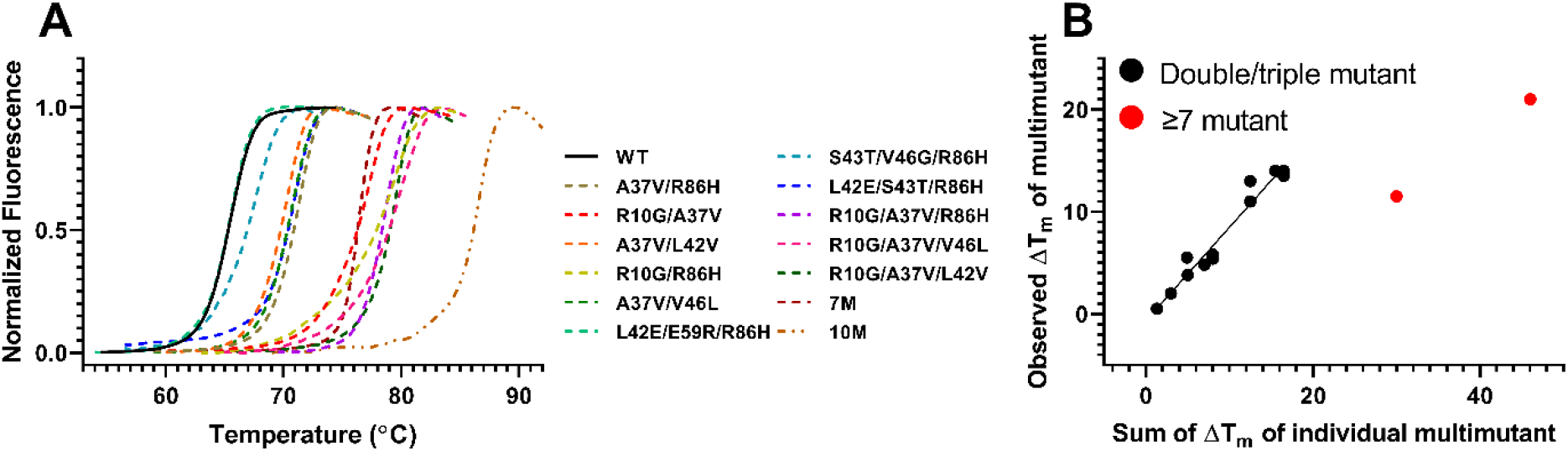
Thermal shift assay data for select CcdB double, triple and multi-site mutants. (A) Data for WT and mutants are shown in black and colour respectively. Multi-mutants showed higher thermal stability compared to individual mutants. Mutations present in 7M are Y8D/R10G/E11P/S12G/A37V/R40S/A69R and 10M are Y8D/R10G/E11P/S12G/A37V/R40S /L42V/V46L/A69R/R86H. (B) The multi-mutants showed an additive effect when two or three mutations were combined. Multi-mutants which contain seven or more mutations did not show completely additive stabilization.

### Thermal aggregation analysis of stabilized mutants

To ascertain the ability of mutations to prevent loss of function from transient high temperature exposure, CcdB mutants were incubated at elevated temperatures for 1 hour, this can result in unfolding of CcdB and subsequent irreversible aggregation (41). The fraction of active protein remaining after incubation was assessed by its ability to bind GyrA at room temperature using SPR. In the case of WT, we did not observe a large decrease in the fraction of active protein except when incubated at temperatures at and above 60 ^º^C (Figure 9A). After incubation at 80 ^º^C, the protein was completely denatured, and did not show any binding with Gyrase. In contrast, one stabilized single mutant R10G and three multi-site mutants retained significant activity after incubation at 80 ^º^C for one hour (Figure 9B). Six mutants showed a higher fraction of active protein than WT after heating at 60 ^º^C (Figure 9C). Three single mutants and two double mutants showed no reduction in active fraction of protein. Surprisingly, the triple mutant, (R10G/A37V/R86H) which showed higher thermal stability than the R10G single mutant, showed higher aggregation at 80 ^º^C, unlike R10G which was partially resistant to aggregation under these conditions. This shows that thermal stability and thermal tolerance need not always be correlated, and stability enhancing and aggregation preventing mutants and mutant combinations can be different.

**Figure 9:**
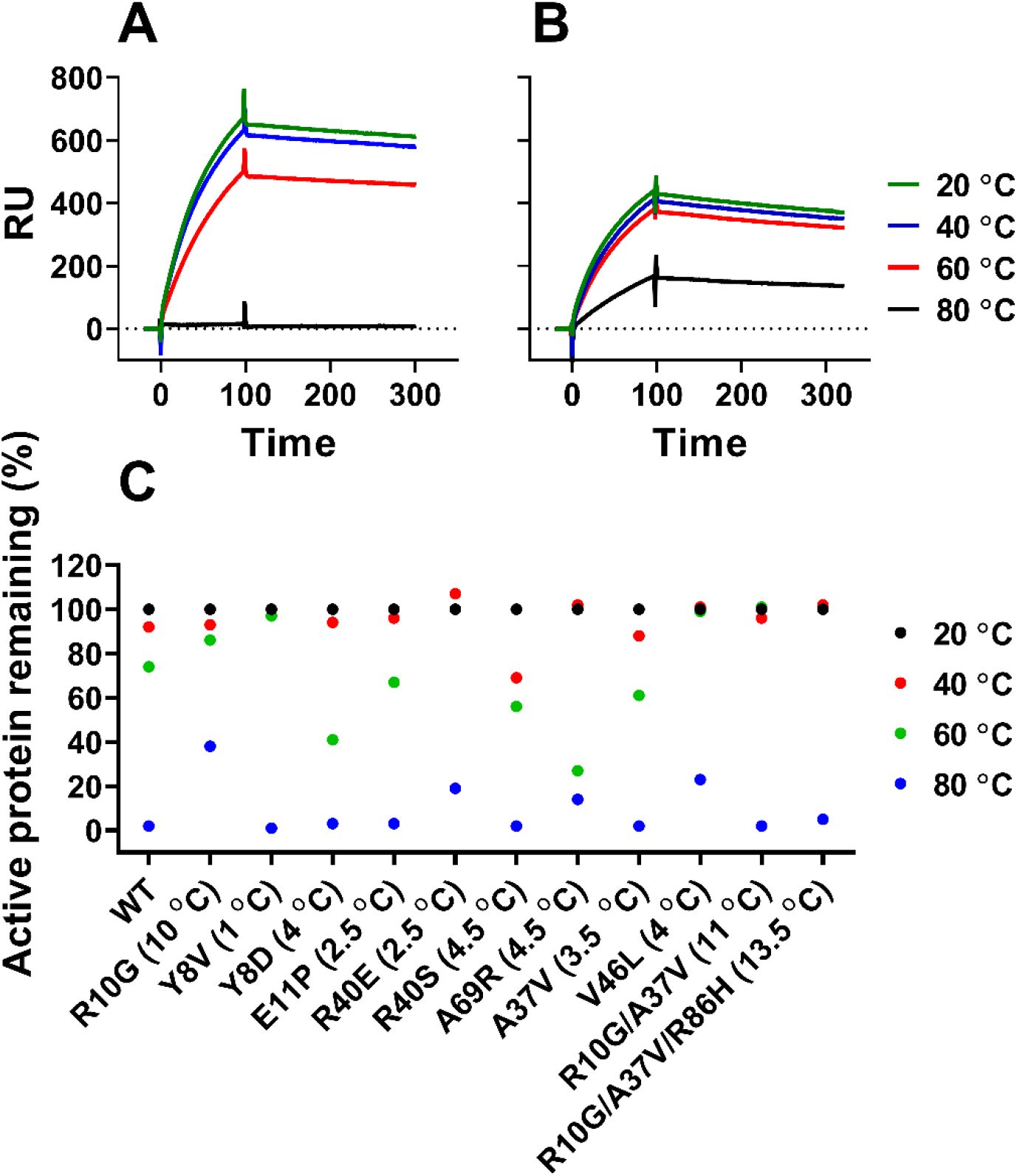
Estimation of the fraction of active protein after thermal stress. CcdB WT and mutants were incubated at 20, 40, 60 and 80 ^º^C for one hour. The fraction of active protein was subsequently estimated by assaying binding to GyrA at 25 ^º^C using SPR. A representative SPR sensorgram for (A) WT CcdB and (B) R10G CcdB showing the relative amount of active protein remaining in the samples after incubation at different temperatures for one hour. (C) Fraction (%) of active protein after incubation at indicated temperature for 1 hour. ΔT_m_ of stabilized mutants is mentioned next to the key of each mutant.

### Sorting of multiple libraries simultaneously

When the suppressor mutations were identified using both enhanced binding to ligand relative to the corresponding PIM, as well as alleviating the destabilizing effect of at least two PIMs as the criteria, predictions were highly specific. Hence, to find a larger number of stabilizing mutants, it is desirable to screen multiple PIM libraries. However, screening multiple individual libraries is laborious. In most FACS based library screens, individual libraries are sorted for multiple rounds to enrich for the best binders. If this approach is applied to pooled libraries with multiple PIMs, the most stable PIM will dominate, resulting in the enrichment of mutants only from this library. To confirm this, we therefore sorted the pooled library as explained (Figure 10A) and we found that mutants with the highest binding were largely from the L36A library as this PIM has the weakest effect on binding (Figure 10B). To overcome this problem, we instead sorted multiple populations based on binding of the pooled library (Figure 10C) and compared the relative binding of putative (PIM, suppressor) pairs with those of individual PIMs. We found a good correlation between the MFI_seq_ (bind) of mutants from the pooled library, with those from individually analysed libraries (Figure 10D), with an increase in the correlation as the read cut-off increased. This suggests that single round sorting of YSD pooled SSSM libraries can rapidly identify stabilized mutations.

**Figure 10:**
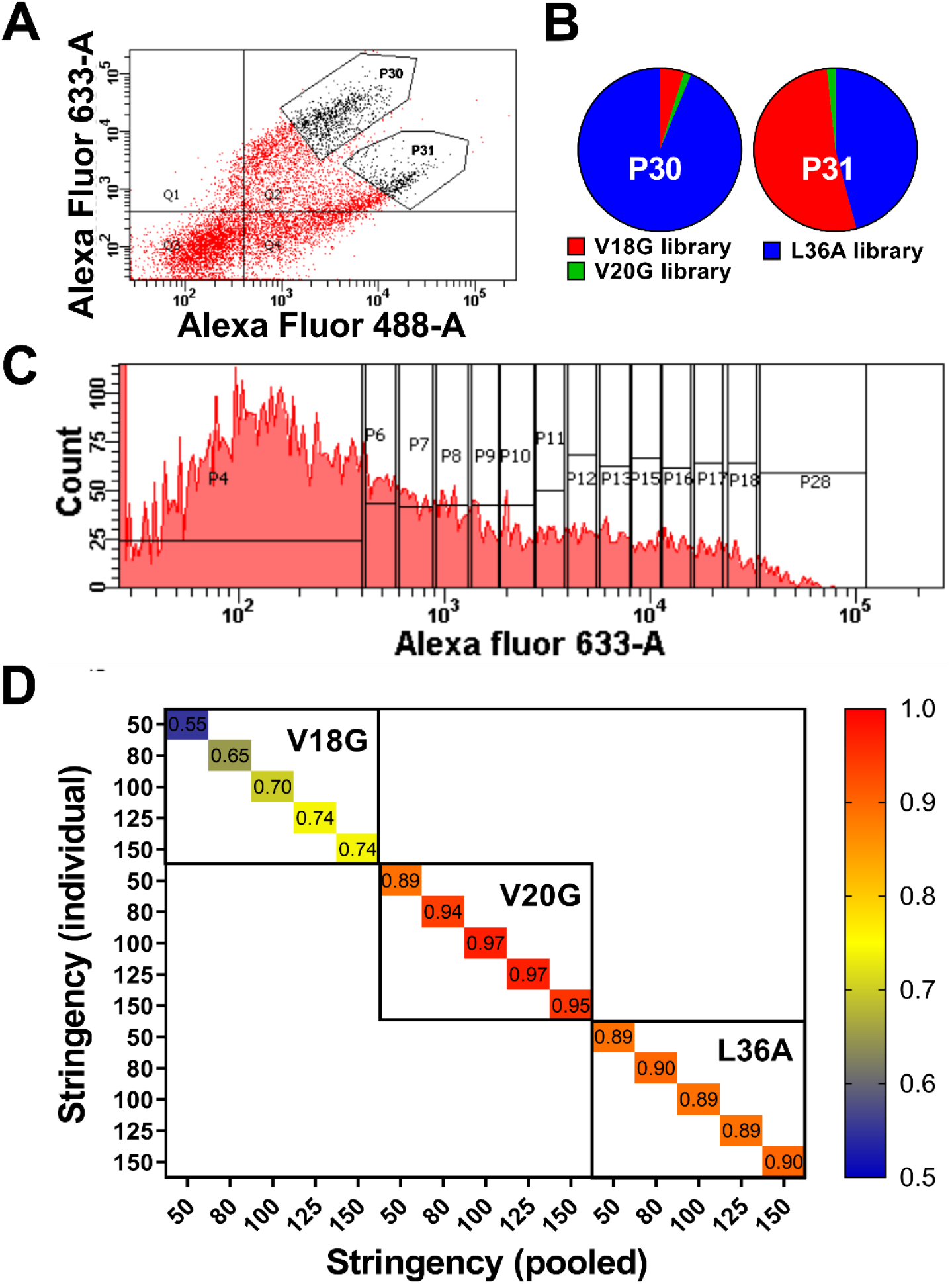
FACS of pooled libraries. (A) Dot plot showing the expression and binding of pooled library. Two different gates, P30 and P31 were used to sort the populations showing the highest expression and binding. (B) Pie chart of relative enrichment of mutants from each library after one round of sorting and deep sequencing in gates P30 and P31. (C) Sorting of pooled V18G, V20G and L36A library based on binding to GyrA14. (D) Heat map of correlation coefficient between the binding MFI of mutants calculated from an individual library and the pooled library at different stringencies, where stringency is the minimum number of reads per mutant.

## Discussion

Directed evolution has drastically reduced the time required to design proteins with new activities. Directed evolution is performed most often in conjunction with display techniques like phage and yeast surface display, which involve selection of better binders to a target of interest from large libraries. While this approach readily selects for high affinity binders, selecting for stable proteins is more difficult. Phage display libraries have very high diversity (42). However, in phage display experiments, there is less control over the selection of populations during enrichment and effects of post-translational modification including glycosylation cannot be studied. Yeast display enables the selection of populations during enrichment using FACS (5). In a recent report, expression level following YSD was used to identify a few stabilized mutants of SARS CoV-2 RBD (39). In the present study, we did not see a good correlation between expression level of individual mutants and thermal stability of stabilized mutants. Instead, we observed that MFI_seq_ (bind) is a better predictor of protein stability than MFI_seq_ (expr). Previously, we found that for stable proteins it is difficult to isolate stabilizing mutants using yeast surface display, as the surface expression and binding of all the mutants above a threshold stability are often similar (22). To overcome this problem, we introduced a PIM in the DMS library of CcdB which reduced binding of all the mutants present in the library and then selected for suppressors which showed improved binding compared to the PIM. We hypothesized that if a stabilizing mutation alleviates the destabilizing effect of at least two PIMs, it is likely to be a true global suppressor. We therefore introduced four different PIMs with varied stability in the DMS library.

We also compared our stabilized mutant prediction with the *in silico* tools DeepDDG (34), PremPS (35), PopMuSiC (36), INPS-MD (37) and PROSS (43). DeepDDG, PremPS, PopMuSiC and INPS-MD did not provide a good estimation of the stability of the stabilized mutants. PROSS is an alternative approach, which uses consensus sequence analysis combined with ROSETTA energy calculations, to predict stabilized protein sequences which contain a large number of (possibly) small effect mutations. PROSS necessarily requires a large number of sequences in a multiple sequence alignment. Also, for some applications, notably in vaccine immunogen design, it is desirable to achieve stabilization through a minimal number of substitutions so as to minimize unwanted changes in surface amino acids that might negatively impact immunogenicity, which is different from and complementary to the PROSS approach.

For libraries with the V18G, V20G and L36A PIMs used in the study, a high specificity for the prediction was observed. These PIM libraries also showed a good correlation between protein stability and MFI_seq_ (bind) (Figure 2). The identified mutants also showed additive stabilization when double or triple mutants were combined.

We also found that if a greater number of PIM libraries are screened, it enhances the prediction of stabilized mutants. To decrease the time and effort involved in screening multiple libraries, we pooled multiple libraries. The pooled library showed similar reconstructed MFIs of mutants to those obtained from individually analysed libraries, validating the approach used. Importantly, the size of libraries employed here is relatively small, identical to deep mutational scanning libraries, with a total size of 32N, where N is the length of the protein sequence. It can thus easily be extended to other library technologies including lentiviral and transposon libraries, phage and mammalian display.

The methodology has the following limitations. The methodology requires a conformation specific ligand to differentiate between the destabilized PIM and stabilized PIM-suppressor pair. For some proteins hyper glycosylation on the yeast cell surface may interfere with the binding between yeast surface displayed protein and its ligand. As an alternative to ligand binding, protease resistance can also be potentially used to select for stabilized mutants (44, 45). However, the aggregation and unfolding on the yeast cell surface may limit the cleavage of such proteins. Use of a protease assay to screen for stabilized mutants, assumes that protease cleavage occurs primarily in the unfolded state (46). While this is true for small well folded proteins, for larger proteins, initial sites of cleavage may occur at surface loops, complicating interpretation of protease screening results. Combing individual mutants from deep mutational scans has limitations and that there are often trade offs between enhancements in stability and binding (47). However, by combining putative stabilizing mutations which are not close to each other or active-site residues we believe that significant stabilization is possible without negatively impacting binding affinity as we have demonstrated both for CcdB in the present work and for the RBD for SARS-CoV-2 in another study (38). Overall the present methodology offers a robust pathway to identify stabilizing mutations in any protein of interest for which a surface display based binding screen is available.

## Experimental Procedures

### Bacterial strains, yeast strains and plasmids

*E*.*coli* strain Top10Gyrase has a mutation in the gyrA gene which prevent CcdB toxicity. The EBY100 strain of *Saccharomyces cerevisiae* has TRP1 mutation which make it auxotrophic for tryptophan and transformants can be selected on minimal media. WT and mutant *ccdB* genes were coned in pBAD24 plasmid for controllable expression in *E*.*coli*. pPNLS shuttle vector was used to display ccdB mutants for yeast cell surface expression.

### Purification of WT and mutant CcdB proteins

CcdB WT and mutant proteins were purified as described (48), briefly, overnight grown culture was diluted 100 folds in 300 ml LB media containing ampicillin (100 µg/ml). The cells were grown and induce at an OD_600_ ∼0.5 for 3 hours at 37 ^º^C. The cells were harvested after induction and lysed using sonication in lysis buffer (10 mM HEPES, 1 mM EDTA, 10% glycerol, pH 8). The soluble fraction was separated using centrifugation and incubated with Affigel-15 coupled to CcdA peptide (residue 45-72) for 2 hours at 4 ^º^C. The unbound fraction was removed, and beads were washed with bicarbonate buffer (50 mM NaHCO3 and 500 mM NaCl). The proteins were eluted with glycine (200 mM, pH 2.5) and collected in tubes containing an equal volume of HEPES buffer (400 mM, pH 8).

### Second site Saturation Suppressor Mutagenesis (SSSM) library generation

A ccdB Deep Mutational Scanning (DMS) library in which each individual residue was randomized, was generated through an inverse PCR based approach (49). Briefly, for a given site, the forward primer had NNK at the 5’end of the primer and the reverse primer starts at the -1 site, relative to the mutation. Individual single-site mutants were generated by inverse PCR, pooled in equimolar ratio, gel extracted, phosphorylated and blunt end ligated at 15 °C. The ligated product was column purified and transformed in electrocompetent *E*.*coli* Top10Gyrase cells. Transformed colonies were scraped and pooled plasmids were purified. The second site suppressor mutant library for CcdB was generated by introducing PIMs in the DMS library as described (29). Briefly, for a given PIM introduction, WT ccdB and ccdB library were amplified in two fragments using two sets of oligos. For each fragment one of the oligos binds to the vector and the other binds to the gene. The primer of both fragments which bind to the gene were completely overlapping and contained the desired PIM mutation. Two separate transformations were performed in yeast (50), in the first transformation, fragment 1 of the ccdB library was combined with fragment 2 of WT ccdB and in the second transformation, fragment 2 of the ccdB library was combined with fragment 1 of WT ccdB. The transformed cells were scraped and pooled, based on the number of mutants present in each transformed plate in an attempt to ensure equal representation of all (PIM, mutant) pairs in the resulting library.

### Yeast surface expression and sorting of SSSM library

Yeast surface display and flowcytometric analysis were performed as explained earlier (22). Briefly, *Saccharomyces cerevisiae* EBY100 cells containing WT ccdB or mutant plasmids were grown in SDCAA (glucose 20g/L, yeast nitrogen base 6.7g/L, casamino acid 5g/L, citrate 4.3g/L, sodium citrate dihydrate 14.3g/L) media for 16 hours and induced in SGCAA (galactose 20g/L, yeast nitrogen base 6.7g/L, casamino acid 5g/L, citrate 4.3g/L, sodium citrate dihydrate 14.3g/L) media for an additional 16 hours at 30 ^º^C. Ten million cells were taken for FACS sample preparation. The second site suppressor mutant (SSSM) library of CcdB was sorted based on 1D sorting of surface expression and binding. The cells were incubated with 200 µl chicken anti-HA antibodies (Bethyl labs, 1:600 dilution) followed by incubation with 200 µl goat anti-chicken antibodies (Invitrogen, 1:300 dilution) conjugated with Alexa fluor 488 to sort the cells based on the cell surface expression. The induced cells were incubated with 200 µl FLAG tagged GyrA14 (1000 nM). The estimated K_d_ for WT CcdB to GyrA14 is 4 nM. A higher CcdB concentration was employed to ensure that even destabilized mutants where only a small fraction was properly folded, showed detectable binding to CcdB. The cells were washed and incubated with 200 µl mouse anti-FLAG antibodies (Sigma, 1:300 dilution), followed by incubation with 200 µl rabbit anti-mouse antibodies (Invitrogen, 1:1500 dilution) conjugated with Alexa fluor 633 to sort the cells based on binding of displayed CcdB mutant to cognate ligand, GyrA14 (29). The sorting of CcdB libraries was performed using a BD Aria III cell sorter. In the case of simultaneous sorting of multiple SSSM libraries, each library samples were prepared separately as explained above and pooled before sorting.

### Sample preparation for deep sequencing

Deep sequencing samples were prepared as explained earlier (22). Briefly, sorted cells were grown on SDCAA agar plates, colonies were scraped, and pooled plasmids were extracted. The ccdB gene was PCR amplified using the primers having multiplex identifier (MID) sequence at the 5’end, that bind upstream and downstream of the ccdB gene to segregate the reads from different sorted bins. The DNA was PCR amplified for 15 cycles, equal amounts of DNA from each sorted population were pooled, gel extracted, and the library was generated using TruSeq™ DNA PCR-Free kit from Illumina. The sequencing was done on an Illumina HiSeq 2500 250PE platform at Macrogen, South Korea.

### Analysis of deep sequencing data

Deep sequencing data for the ccdB mutants were processed as described (22). Briefly, the paired end reads were assembled using the PEAR v0.9.6 (Paired-End Read Merger) tool (51). Following assembly, reads were filtered to eliminate those that do not contain the relevant MID and/or primers along with the reads having mismatched MID’s. Only those reads which have bases with Phred score ≥ 20 are retained. Reads which pass the assembling and filtering step were binned according to the respective MIDs. Binned reads were aligned with the wild type ccdB sequence using the Water v6.4.0.0 program (52) and reformatted. Finally the reads were classified based on insertions, deletions and substitutions (single, double, etc mutants).

### MFI reconstruction from deep sequencing data

The MFI of each mutant was reconstructed as described (22). Briefly, reads of each mutant were normalized across different bins (Equation 1). The fraction of each mutant (X*i*) distributed across each bin was calculated (Equation 2). The reconstructed MFI (MFI_seq_) of individual mutant was calculated by the summation of the product, obtained upon multiplying the fraction (X*i*) of that mutant in bin (*i*) with the MFI of the corresponding bin obtained from the FACS experiment (F*i*), across the various bins populated by that mutant (Equation 3). The normalized MFI of each mutant was calculated from the reconstructed MFI of each mutant (Equation 4).

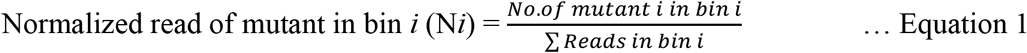

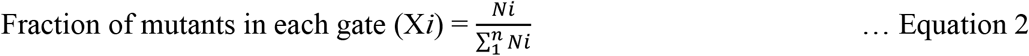

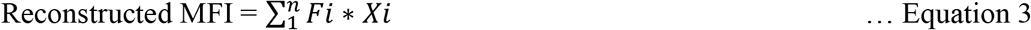

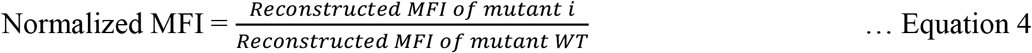

MFI_seq_ (expr) and MFI_seq_ (bind) refer to reconstructed values from FACS sorting based on mutant expression and binding to GyrA14 respectively. Stabilized mutants were classified as those which showed at least 25% enhanced binding or expression when present as PIM-suppressor pair compared to the PIM alone.

### Protein thermal stability measurement

This was carried out as described (48). Briefly, a solution of total volume 20 μL containing 10 μM of the purified CcdB protein and 2.5x Sypro orange dye in buffer (200 mM HEPES, 100 mM glycine), pH 7.5 was heated from 15 ^º^C to 90 ^º^C with 0.5 ^º^C increment every 30 seconds on an iCycle iQ5 Real Time Detection System (Bio-Rad, Hercules, CA). The normalized fluorescence data was plotted against temperature (53).

### Thermal aggregation studies of CcdB mutants

CcdB mutants and WT proteins (500 nM, 200 µL for each of the proteins) were subjected to incubation at four different temperatures, 20, 40, 60 and 80 ^º^C on an iCycle iQ5 Real Time Detection System

(Bio-Rad, Hercules, CA). The temperature was gradually increased to the desired temperature at a rate of 3 ^º^C/min and samples were kept at the desired temperature for 1 hour. The heated protein was then cooled down to 4 ^º^C at the rate of 3 ^º^C/min. The aggregated protein was removed using centrifugation at 18000g. The fraction of active protein remaining was measured by binding to GyrA14 on a Biacore 2000 SPR platform. The percentage of active protein at different temperatures was calculated using the following equation.

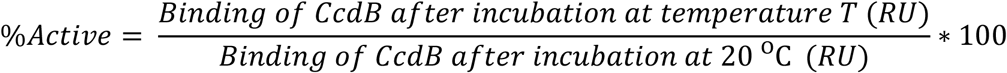

## Data Availability

The deep sequencing data discussed in the present study has been deposited in NCBI’s Sequence Read Archive (accession no. SRR16094780). Illumina sequencing counts for each ccdB double mutant of FACS bins are available at https://github.com/rvaradarajanlab/ccdb_sssm/blob/main/ccdn_sssm_freq.xlsx. MFI_seq_ (expr) and MFI_seq_ (bind) of CcdB mutants are available at https://github.com/rvaradarajanlab/ccdb_sssm/blob/main/Supplementary_data_sssm_calc_MFI.xlsx. Remaining data is available in the manuscript.

## Supporting information

This article contains supporting information.

## Acknowledgements

S.A. acknowledges Council of Scientific & Industrial Research for his fellowship (SPM-07/079(0218)/2015-EMR-I). K.M. is thankful to Department of Science and Technology (DST) Science and Engineering Research Board for financial support, sanction order no: PDF/2017/002641. Aparna Asok is duly acknowledged for FACS. Nonavinakere Seetharam Srilatha is duly acknowledged for the SPR experiments. Munmun Bhasin is acknowledged for deep sequencing data submission at NCBI’s Sequence Read Archive.

## Author contribution

R.V. and S.A. designed the experiments. S.A. performed all the experiments. R.V. and S.A. analyzed all the data. K.M. wrote the software and carried out the processing of the deep sequencing data. S.A. analyzed the deep sequencing data of CcdB. G.C. performed SPR to estimate the dissociation constant of CcdB mutants. R.V. and S.A. wrote most of the manuscript.

## Funding and additional information

This work was funded by grants to RV from the Department of Science and Technology, grant number-EMR/2017/004054, DT.15/12/2018), Government of India, Department of Biotechnology, grant no. BT/COE/34/SP15219/2015 DT. 20/11/2015, Ministry of Science and Technology, Government of India and Bill and Melinda Gates Foundation (USA) (INV-005948). We also acknowledge funding for infrastructural support from the following programs of the Government of India: DST FIST, UGC Centre for Advanced study, Ministry of Human Resource Development (MHRD), and the DBT IISc Partnership Program. The funders had no role in study design, data collection and interpretation, or the decision to submit the work for publication.

## Statistical Analysis

All the data were plotted using the GraphPad Prism software 9.0.0. The correlation coefficients between deep sequencing replicates were estimated using the GraphPad Prism software 9.0.

## Abbreviations

YSD: Yeast surface display
DMS: Deep Mutational Scanning
SSSM: Second site Saturation Suppressor Mutagenesis
FACS: Fluorescence-activated cell sorting
PIM: Parent Inactive Mutation
MID: multiplex identifier

## Supporting Information

**Figure S1:**
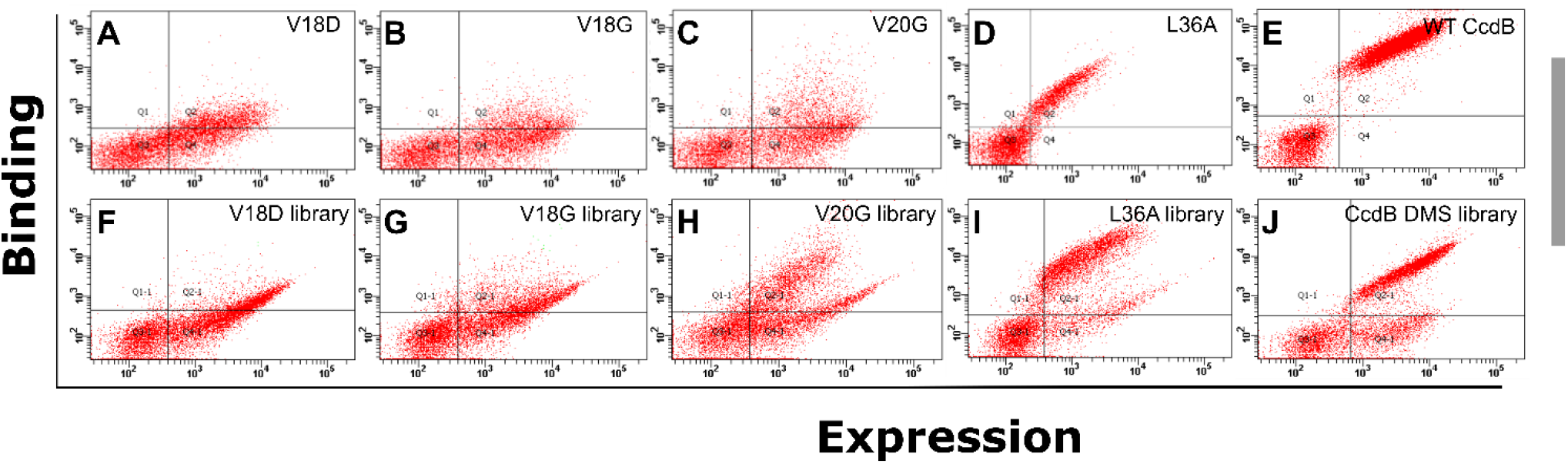
FACS double plot showing the expression and binding to GyrA14 of CcdB PIMs, WT and their corresponding DMS libraries. FACS double plot of CcdB PIMs (A) V18D, (B) V18G, (C) V20G and (D) L36A and (E) WT CcdB. Double plots of DMS libraries in the background of (F) V18D, (G) V18G, (H) V20G (I) L36A (J) WT CcdB

**Figure S2:**
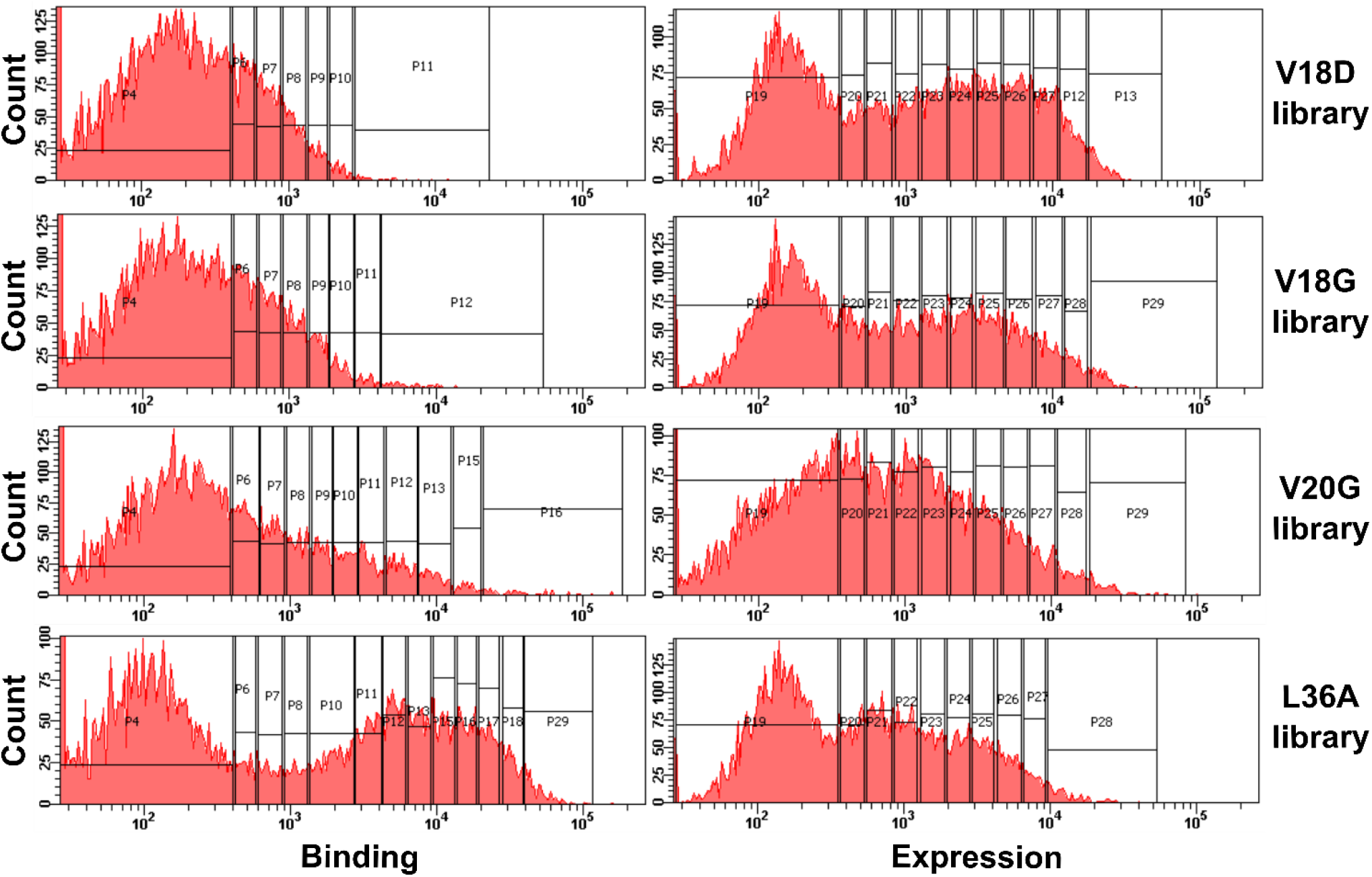
Histograms of expression and binding to GyrA14 of PIM containing CcdB libraries. Left and right panels show binding and expression histograms. Different populations sorted into bins based on expression and binding are indicated by the vertical lines.

**Figure S3:**
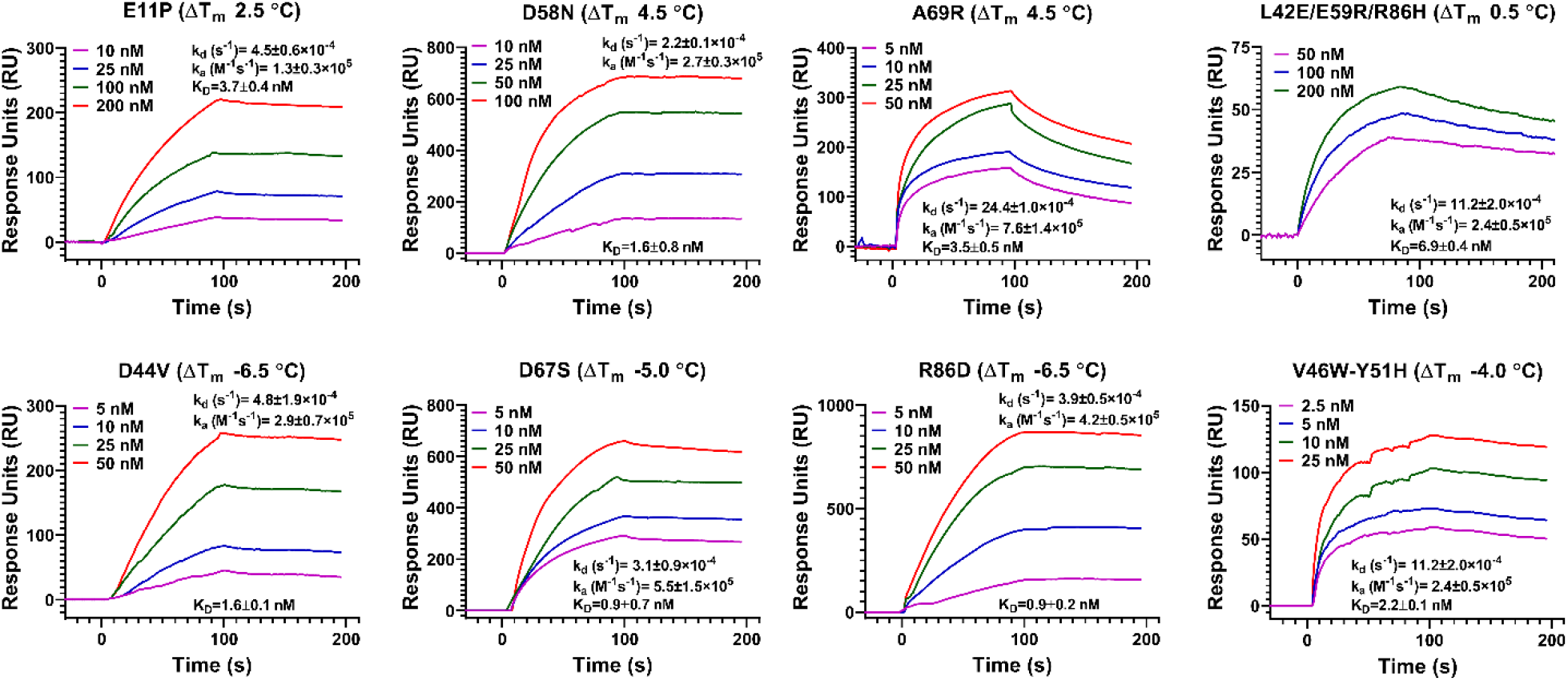
Binding of GyraseA-14 to CcdB mutant proteins. The ligand GyrA14 was immobilized on the CM5 chip by standard amine coupling. Binding was measured by passing varying concentrations of the analyte (CcdB proteins) over the ligand (GyrA14) immobilised chip. Overlays show the binding kinetics of ligand with different concentrations of analyte. Top panels shows the binding kinetics of stabilized single or multi-mutants, bottom panel shows the binding kinetics of destabilized single or multi-mutants. The data was fitted to the 1:1 Langmuir interaction model to obtain the kinetic parameters.

**Table S1:** Number of mutants analysed in each PIM library for expression and binding.

|  |  | **Binding** | **Expression** |
| --- | --- | --- | --- |
| **V18D library** | **Codon level** | **1275** | **1139** |
|  | **Codon averaged** | **935** | **862** |
| **V18G library** | **Codon level** | **1319** | **1353** |
|  | **Codon averaged** | **967** | **989** |
| **V20G library** | **Codon level** | **1237** | **1343** |
|  | **Codon averaged** | **890** | **977** |
| **L36A library** | **Codon level** | **1763** | **670** |
|  | **Codon averaged** | **1189** | **530** |

**Table S2:**
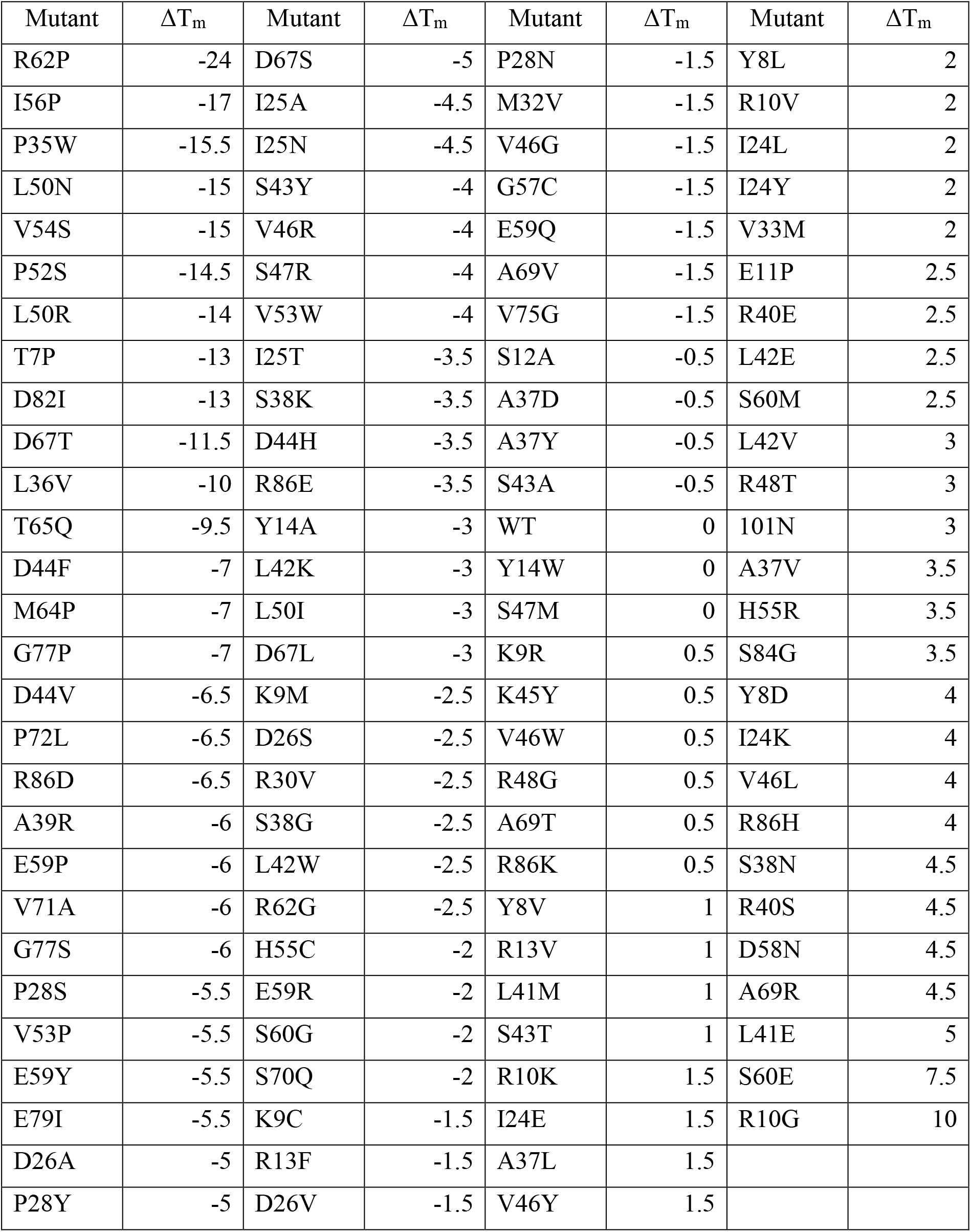
Thermal stability of CcdB mutants estimated using TSA.

**Table S3:** PROSS predictions of CcdB stabilizing mutations. For three designs PROSSS did not predict any mutation, for the remaining designs 2-16 mutations combinations were predicted. The mutations predicted by PROSS are highlighted with different colours where, grey colour indicates the mutants for which data is not available by saturation suppressor mutagenesis. Orange colour indicate active-site mutants, green colour indicates predicted stabilizing mutation both by saturation suppressor mutagenesis and PROSS, red colour indicates destabilizing mutation predicted by saturation suppressor mutagenesis and stabilizing mutation predicted by PROSS.

| Residue | Wild Type | Design 1 | Design 2 | Design 3 | Design 4 | Design 5 | Design 6 | Design 7 | Design 8 | Design 9 |
| --- | --- | --- | --- | --- | --- | --- | --- | --- | --- | --- |
| 49 | E |  |  |  |  |  |  |  |  | R |
| 24 | I |  |  |  |  |  |  |  |  | L |
| 92 | N |  |  |  |  |  |  |  |  | A |
| 13 | R |  |  |  |  |  |  |  |  | A |
| 47 | S |  |  |  |  |  |  |  |  | P |
| 58 | D |  |  |  |  |  |  |  | G | G |
| 87 | E |  |  |  |  |  |  |  | R | R |
| 57 | G |  |  |  |  |  |  |  | N | N |
| 15 | R |  |  |  |  |  |  |  | K | K |
| 37 | A |  |  |  |  |  |  | V | I | F |
| 88 | N |  |  |  |  |  |  | D | D | D |
| 95 | N |  |  |  |  |  |  | D | D | D |
| 41 | L |  |  |  |  | H | H | H | H | H |
| 60 | S |  |  |  |  | D | D | D | D | D |
| 38 | S |  |  |  | P | P | P | P | P | P |
| 61 | W |  |  |  | Y | Y | Y | Y | Y | Y |

